# MathFeature: Feature Extraction Package for Biological Sequences Based on Mathematical Descriptors

**DOI:** 10.1101/2020.12.19.423610

**Authors:** Robson P. Bonidia, Danilo S. Sanches, André C.P.L.F. de Carvalho

## Abstract

Machine learning algorithms have been very successfully applied to extract new and relevant knowledge from biological sequences. However, the predictive performance of these algorithms is largely affected by how the sequences are represented. Thereby, the main challenge is how to numerically represent a biological sequence in a numeric vector with an efficient mathematical expression. Several feature extraction techniques have been proposed for biological sequences, where most of them are available in feature extraction packages. However, there are relevant approaches that are not available in existing packages, techniques based on mathematical descriptors, e.g., Fourier, entropy, and graphs. Therefore, this paper presents a new package, named MathFeature, which implements mathematical descriptors able to extract relevant information from biological sequences. MathFeature provides 20 approaches based on several studies found in the literature, e.g., multiple numeric mappings, genomic signal processing, chaos game theory, entropy, and complex networks. MathFeature also allows the extraction of alternative features, complementing the existing packages.

**Availability and implementation:** MathFeature is freely available at https://bonidia.github.io/MathFeature/ or https://github.com/Bonidia/MathFeature

## 1 Background

In the last years, Machine learning (ML)-based tools have been developed for various genomics, transcriptomics, and proteomics problems [1]. Nevertheless, for the successful application of ML algorithms, relevant features need to be extracted, to represent the main aspects of the original sequence. In [2, 3], the authors address the relevance of using an appropriate mathematical expression to extract features from biological sequence data. Based on this, many techniques have been developed and investigated to extract numerical representative information from sequences [4, 5]. In which, the main challenge is how to numerically represent a biological sequence in a numeric vector. To deal with this challenge, several of these features were made available in public software packages, such as: PseKNC-General [5], PseKNC [4], Pse-in-One [3], DNAshapeR [6], repDNA [7], repRNA [8], Pse-in-One 2.0 [9], BioSeq-Analysis [10], iFeature [11], PyFeat [12], Seq2Feature [13], iLearn [14].

These previous studies have produced tools, packages, web servers, or toolkits. However, each of them was limited regarding the mathematical features made available (e.g., multiple numeric mappings, Fourier, chaos game theory, entropy, and complex networks). Therefore, in this work, we present an open-source Python package, named MathFeature, which provides in a single environment, many of the mathematical features previously proposed for feature extraction from biological sequences. MathFeature provides 20 mathematical approaches, here named descriptors, organized into five categories. To our best knowledge, MathFeature is the first package to provide such a large and comprehensive set of mathematical feature extraction descriptors for biological sequences.

## 2 Package Description

In MathFeature, the descriptors are applied to a sequence according to the pipelines illustrated by Figure 1. In Table 1, we organize the 20 descriptors into 5 groups, according to how they work. Furthermore, we have developed a user-friendly tool that covers several mathematical descriptors. Next, we also classified the main aspects of each group, as follow:

- **Numerical Mapping:** Several sequence analysis studies require converting a biological sequence to a numeric sequence. Previous studies have proposed different descriptors for such, which are able to represent important aspects of these sequences. This group contains 7 descriptors for numerical mapping: Voss [15] (known as binary mapping), Integer [16], Real [17], Z-curve [18], EIIP [19], Complex Numbers [20, 21] and Atomic Number [22, 23].
- **Chaos Game Representation (CGR):** This approach is also a mapping, but scale-independent and iterative for geometric representation of DNA sequences [24]. There are several variations of this group, in this package, we included classical CGR [24, 25], frequency CGR [26], and CGR signal with Fourier Transform (FT) [25].
- **Fourier Transform:** This group extracts sequence features based on Genomic Signal Processing, using FT, a widely applied approach in several biological sequence analysis problems [27, 25, 23, 28]. To implement GSP techniques, we use all numerical mappings. A mathematical exploration can be seen in [28].
- **Entropy:** Various studies have applied concepts from information theory for sequence feature extraction, mainly Shannon’s entropy [29, 30]. However, according to [31], another entropy-based measure has been successfully explored in several studies, Tsallis entropy [32], proposed to generalize the Boltzmann/Gibbs’s traditional entropy. This group includes these two descriptors [28].
- **Graphs:** This group has descriptors based on graph theory, a currently very active research area, with relevant results in biology sequences [33, 34]. In addition, the descriptors included in this group were proposed in [35] and also explored in [28].

**Table 1:** Descriptors calculated by MathFeature for DNA, RNA, and Protein sequences.

| Descriptor groups | Descriptor | Dimension | Biological Sequence |  |
| --- | --- | --- | --- | --- |
| *Numerical Mapping* | Binary | $L \cdot 4$ | DNA/RNA | $L =$ |
| | Z-curve | $L \cdot 3$ | DNA/RNA | |
| | Real | $L$ | DNA/RNA | |
| | Integer | $L$ | DNA/RNA | |
| | EIIP | $L$ | DNA/RNA | |
| | Complex Number | $L$ | DNA/RNA | |
| | Atomic Number | $L$ | DNA/RNA | |
| *Chaos Game* | Chaos Game Representation | $L \cdot 2$ | DNA/RNA | |
| | Frequency Chaos Game Representation | $L - k + 1$ | DNA/RNA | |
|  | Chaos Game Signal (with Fourier) | 19 | DNA/RNA |  |
| *Fourier Transform* | Numerical Mapping + Fourier | 19 | DNA/RNA |  |
| *Entropy* | Shannon | $k$ | DNA/RNA/Protein | |
| | Tsallis | $k$ | DNA/RNA/Protein | |
| *Graphs* | Complex Networks (Codons) | $12 \cdot t$ | DNA/RNA/Protein | |
| | Complex Networks (different k-mers) | $12 \cdot t \cdot k$ | DNA/RNA/Protein | |

length of the longest sequence, $k$ = frequencies of k-mer, $t$ = threshold - number of subgraphs.

**Figure 1:**
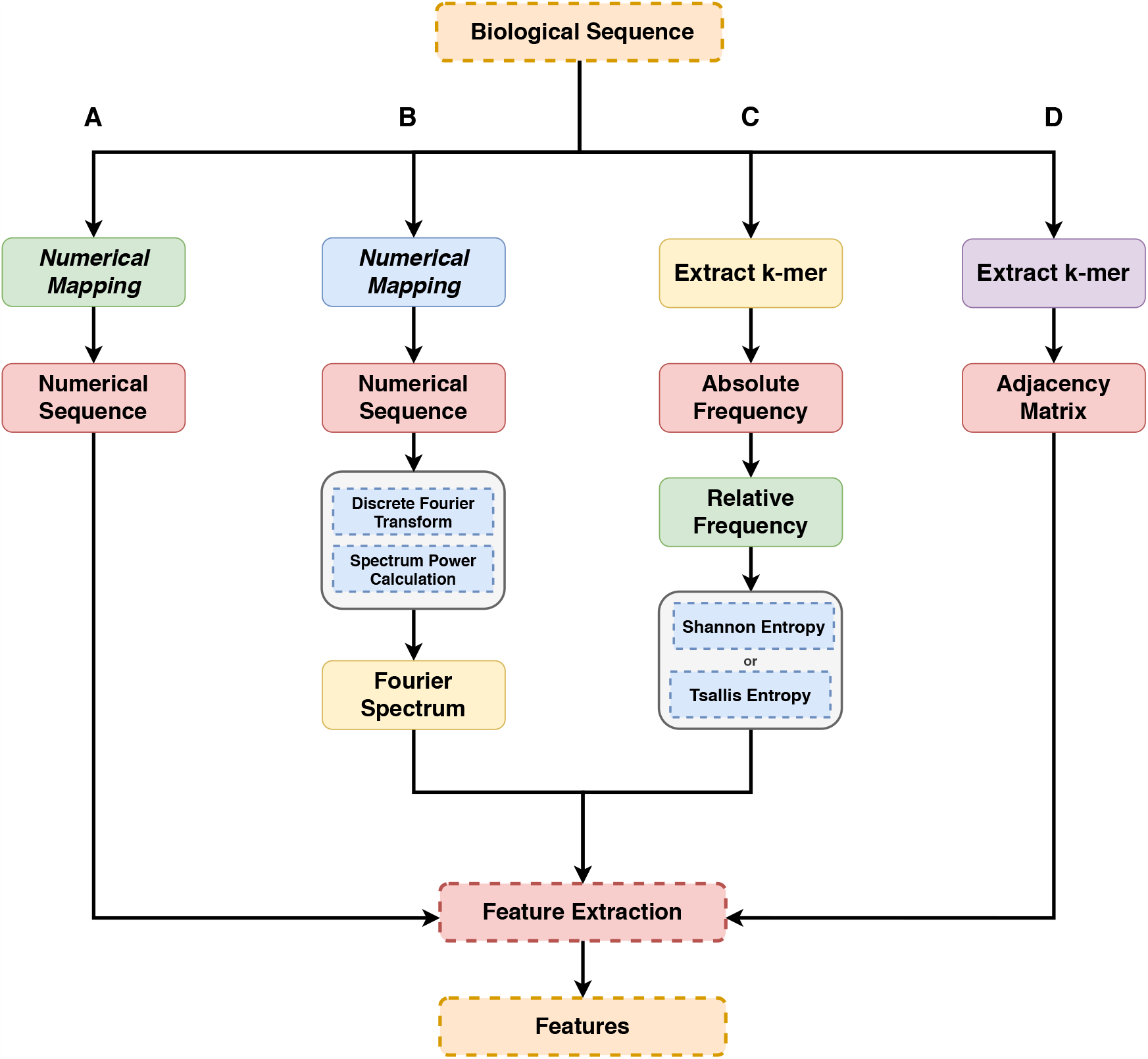
Pipeline of descriptors calculated by MathFeature. **A:** Numerical Mapping and Chaos Game Representation; **B:** Fourier Transform; **C:** Entropy; **D:** Complex Networks.

## 3 Results

In this study, we presented MathFeature, a novel feature extraction package based on mathematical features. MathFeature provides 20 descriptors to numerically represent biological sequences, organized in 5 groups: multiple numeric mappings, Fourier transform, chaos game theory, entropy, and complex networks. Our main purpose is to complement the existing feature extraction packages, providing alternatives to extract new and relevant features using mathematical descriptors. These descriptors have been previously applied to biological sequences with relevant and robust results, e.g., [28] (performances (ACC) between 0.8901-0.9606), [25], [27], [30], and [35]. MathFeature is freely available at https://github.com/Bonidia/MathFeature and its documentation is provided at https://bonidia.github.io/MathFeature/.

## Acknowledgements

The authors would like to thank USP and CAPES - Finance Code 001 and PROEX-11919694/D for the financial support for this research. *Conflict of Interest:* none declared.

